# Histone deacetylase inhibition reduces deleterious cytokine release induced by ingenol stimulation

**DOI:** 10.1101/193946

**Authors:** Erin T. Larragoite, Racheal A. Nell, Laura J. Martins, Louis R. Barrows, Vicente Planelles, Adam M. Spivak

## Abstract

**Introduction:** Latency reversal agents (LRAs), such as protein kinase C (PKC) agonists, constitute a promising strategy for exposing and eliminating the HIV-1 latent reservoir. PKC agonists activate NF-κB and, in turn, induce deleterious pro-inflammatory cytokine production. Adjuvant pharmacological agents, such as ruxolitinib, a JAK inhibitor, and rapamycin, an mTOR inhibitor, have previously been combined with LRAs to reduce deleterious pro-inflammatory cytokine secretion without inhibiting HIV-1 viral reactivation in vitro. Histone deacetylase inhibitors (HDACi) are known to dampen pro-inflammatory cytokine secretion in the context of other diseases and can synergize with other LRAs to bring dormant proviruses out of latency. In this study we investigated whether a broad panel of epigenetic modifiers, including HDACi, could effectively dampen PKC-induced pro-inflammatory cytokine secretion during latency reversal.

**Methods:** We screened an epigenetic modifier library to identify compounds that reduced intracellular IL-6 production induced by the PKC agonist Ingenol-3,20-dibenzoate. We further tested the most promising epigenetic inhibitor class, HDACi, for their ability to reduce a broad panel of pro-inflammatory cytokines and reactivate latent HIV-1 *ex vivo*.

**Results:** We identified nine epigenetic modulators that reduced PKC-induced intracellular IL-6. In cells from aviremic individuals living with HIV-1, the HDAC1-3 inhibitor, suberohydroxamic acid (SBHA), reduced secretion of pro-inflammatory cytokines TNF-α, IL-5, IL-2r, and IL-17 but did not significantly reactivate latent HIV-1 when used in combination with Ingenol-3,20-dibenzoate.

**Conclusion:** The addition of SBHA to Ingenol-3,20-dibenzoate reduces deleterious cytokine production during latency reversal but does not induce significant viral reactivation in aviremic donor PBMCs. The ability of SBHA to reduce PKC-induced pro-inflammatory cytokines when used in combination with Ingenol-3,20-dibenzoate suggests that SBHA can be used to reduced PKC induced pro-inflammatory cytokines but not to achieve latency reversal in the context of HIV-1.

## 1. Introduction

Antiretroviral therapy (ART) has improved the lives of individuals living with HIV-1 by durably blocking viral replication, reducing plasma viremia levels below the limit of detection (∼20-50 copies/ml) [1-3], and allowing for reconstitution of the adaptive immune system. However, ART does not result in HIV-1 eradication due to the presence of a transcriptionally silent, long-lived latent viral reservoir [4-6]. The presence of this latent reservoir makes ART a lifelong necessity [7]. Additionally, not all patients have access to ART or are compliant with ART regimens due to high costs [8], side effects [9], and social stigma [10, 11]. Aviremic individuals taking ART are prone to low-level chronic immune activation and exhaustion [12], cardiovascular disease [13, 14], non-AIDS related cancers [15], and neurologic complications [16], underscoring the importance of strategies targeting the latent HIV-1 reservoir.

The “shock and kill” strategy [17] proposes that a latency reversing agent (LRA) be used to transcriptionally reactivate the latent reservoir followed by targeting infected cells with immune-mediated mechanisms [17, 18]. Immunological strategies include the use of broadly neutralizing antibodies (bNAbs), cytokines, vaccines, chimeric antigen receptors, and other immune-mediated mechanisms to enhance recognition of HIV-1 infected cells [19, 20]. While all strategies hold merit, enhanced targeting of latently infected cells via immune-mediated mechanisms would still be dependent on unmasking the latent reservoir.

Several mechanistic classes have emerged as potential LRAs including positive transcription elongation factor b (PTEFb) activators [21, 22], cytokines [23], histone deacetylase inhibitors (HDACi) [24-28], and protein kinase C (PKC) agonists [18, 28-30]. Spina *et al*. [29] demonstrated that PKC agonists are uniquely efficacious LRAs across multiple *in vitro* models of HIV-1 latency, whereas other well characterized LRAs such as HDACi had functionality only in select *in vitro* models. PKC agonists are thought to reactivate latent HIV-1 though activation of NF-κB and have been shown to additionally activate AP-1 and NFAT [31]. On the other hand, PKC agonists have undesired effects, including induction of T cell activation [32] and pro-inflammatory cytokine secretion. We previously demonstrated that latency reversal with the PKC agonist Ingenol-3,20-dibenzoate (IDB), used in combination with a second pharmacological agent, ruxolitinib (an FDA approved Janus Kinase [JAK] inhibitor), led to potent reactivation of latent HIV-1 in the absence of pro-inflammatory cytokine secretion[33]. Additionally, rapamycin, an inhibitor of mTOR, has been shown to reduce pro-inflammatory cytokine secretion induced upon reactivation of latent HIV-1 with CD3 and CD28 antibodies [34]. These studies indicate that latency reversal and pro-inflammatory cytokine secretion can be uncoupled. In the present study we expanded our search for epigenetic modifiers that could be used as an adjuvant therapy to reduce cytokine secretion induced by PKC agonists.

While ruxolitinib and rapamycin have been shown to reduce *in vitro* cytokine secretion in the context of HIV-1, HDACi have also been observed to inhibit pro-inflammatory cytokine secretion in vivo in the context of graft-versus-host disease [35] and rheumatoid arthritis [36] as well as in vitro, in the context of LPS stimulation of human peripheral blood mononuclear cells (PBMCs) [37]. Additionally, HDACi are well-known HIV-1 LRAs and it has been shown that they can have additive or synergistic activity with PKC agonists when reactivating latent HIV-1 [27, 38, 39]. Therefore, we hypothesized that HDACi and potentially other epigenetic modifiers could decrease PKC-induced pro-inflammatory cytokines secretion and be used as a combination therapy with PKC agonists to reactivate latent HIV-1. In this study, we performed a screen of epigenetic modifiers and report that suberohydroxamic acid (SBHA), an HDAC1- and 3-inhibitor, suppresses PKC-induced pro-inflammatory cytokines but does not significantly reactivate latent HIV-1.

## 2. Materials and Methods

### Participants

Participants living with and without HIV-1 infection were recruited in accordance with active University of Utah Institutional Review Board (IRB) protocols 58246 and 67637 respectively. Participants living with HIV-1 infection were aviremic (plasma viral loads less than 50 HIV-1 RNA copies/mL) for a minimum of 6 months and prescribed an ART regimen initiated during chronic HIV-1 infection for a minimum of 12 months.

### In vitro epigenetic inhibitor screening

PBMCs were isolated from healthy donors using a Lymphoprep density gradient (Cat# 07861, StemCell Technologies) prior to being cultured in RPMI medium supplemented with 10% FBS and 5 U/mL penicillin/streptomycin overnight to remove monocytes via adherence. Non-adherent PBMCs were then cultured at a density of 1 × 10^5^ cells/100µL in the presence of 100nM compounds from a Cayman Chemical Epigenetics Screening Library obtained from the University of Utah Drug Discovery Core Facility (Item No. 11076) or 100nM ruxolitinib (Cayman Chemical Co., Item No. 11609), a control for cytokine inhibition, for 1.5 h at 37°C. Cells were then incubated in medium alone or the presence of 100nM IDB for 40 hours post-stimulant exposure. At 40 hours post-stimulant exposure, 0.067 µL/100µL BD GolgiStop™ Protein Transport Inhibitor (Cat# 554724) was added to each sample to inhibit cytokine secretion. Cells were then fixed and stained at 48 hours prior to flow cytometry analysis.

### Intracellular cytokine staining and flow cytometry

Cells were washed with 1x phosphate buffered saline (PBS) prior to staining the cells with 0.1µL/100 µL Fixable Viability Dye eFluor^®^ 450 (Cat# 65-0863-14, Affymetrix eBioscience) for 30 min at 4°C. Cells were then washed with 1x PBS prior to fixing with 100 µL BD Cytofix/Cytoperm™ for 30 min at 4°C. Once cells were fixed, they were washed with a perm/wash solution (1x PBS, 3% FBS, 0.1% Saponin, 0.05% Sodium Azide) prior to staining cells in 100 µL perm/wash with 0.5 µL APC anti-human IL-6 antibody (Cat# 501112, Biolegend®) overnight at 4°C. Cells were finally washed in 1x PBS and re-suspended in PBS prior to flow cytometry [BD FACSCanto™ flow cytometer with FACSDiva™ acquisition software (Becton-Dickinson, Mountain View, CA)] and analysis with FlowJo (TreeStar Inc, Ashland, OR).

### Selection of compounds

The percentage of IL-6-positive cells in IDB-alone-treated cells was compared to cells treated with epigenetic modifiers and IDB in order to calculate intracellular IL-6 fold-change. The top epigenetic modifiers that reduced intracellular IL-6 by four-fold or greater (‘hits’) were selected for testing in aviremic patient cells to determine whether these compounds would diminish PKC induced pro-inflammatory cytokines.

### Ex vivo cell culture, qPCR, and cytokine measurements

Peripheral blood mononuclear cells (PBMCs) (Lymphoprep™, Cat # 07861, StemCell Technologies) and resting CD4^+^ T cells (rCD4s) (EasySep™ Human Resting CD4^+^ T Cell Isolation Kit, Cat # 17962, StemCell Technologies) were isolated from HIV-1 aviremic donors living with HIV-1. PBMCs were cultured at a density of 3 × 10^6^/ml in RPMI supplemented with 10% FBS and 5U/mL penicillin/streptomycin. Two epigenetic modifiers (SBHA and panobinostat) that decreased intracellular IL-6 to the highest degree in the *in vitro* IL-6 screen were added at 75nM or 100nM concentrations in the presence of medium or IDB for 48 hours to evaluate *ex vivo* cytokine secretion. Cells cultured in medium alone were used as controls. Supernatant was collected and sent to ARUP Laboratories to measure *ex vivo* cytokine secretion via a commercially available quantitative multiplex bead assay to measure the following cytokines: interferon gamma (IFN-γ), tumor necrosis factor alpha (TNF-α), interleukin (IL) 1 beta (IL-1β), IL-2, soluble IL-2 receptor (IL-2r), IL-4, IL-5, IL-6, IL-8, IL-10, IL-12, and IL-13.

Resting CD4s were cultured at a density of 5 × 10^6^/mL in RPMI supplemented with 10% FBS and 5U/mL penicillin/streptomycin. Epigenetic modifiers (chidamide, apicidin, entinostat, suberohydroxamic acid (SBHA), pyroxamide, panobinostat, WDR5-0103, iniparib and CCG-100602) that reduced intracellular IL-6 were added at 50nM to rCD4s alone or in combination with IDB (45nM) for 48 hours. Medium or DMSO and Dynabead™ Human T-Activator αCD3/αCD28 (αCD3/αCD28) (Cat# 11132D, Gibco™) stimulated cells were used as controls. At 48 hours post-stimulation, supernatant was collected and the <u>r</u>apid *<u>e</u>x <u>v</u>ivo* <u>e</u>valuation of <u>a</u>nti-<u>l</u>atency assay (REVEAL) assay was performed as previously described [28] to quantify viral release.

### Statistical Analysis

Significant changes in reactivation or the secretion of pro-inflammatory cytokines in cells treated with epigenetic modifiers in combination with IDB compared to IDB alone was calculated with GraphPad Prism Version 5.0f (GraphPad Software, Inc., San Diego CA). Statistical significance was calculated using a non-parametric Wilcoxon matched-pairs signed rank test.

## 3. Results

HDACi have been previously described to inhibit cytokine secretion [35-37]. Therefore, we investigated the ability of HDACi and other epigenetic modifiers to reduce pro-inflammatory cytokine secretion upon latency reversal with a PKC agonist. We conducted a screen of 96 epigenetic modifiers (Cayman Chemical Co.) in primary CD4^+^ T cells to identify those that more efficiently quelled the induction of pro-inflammatory cytokines during latency reversal with ingenol-3,20-dibenzoate (IDB). We used IL-6 as a prototypical cytokine for the screen. Nine epigenetic modifiers were identified which reduced intracellular IL-6 levels by four-fold or greater in the context of IDB stimulation (**Figure 1**). These ‘hits’ include six HDACi (Chemical Abstracts Service (CAS) # 743420-02-2 (chidamide), CAS # 183506-66-3 (apicidin), CAS # 209783-80-2 (entinostat), CAS # 38937-66-5 (suberohydroxamic acid (SBHA)), CAS # 382180-17-8 (pyroxamide), CAS # 404950-80-7 (panobinostat)), one mixed lineage leukemia (MLL) inhibitor Cas # 890190-22-4 (WDR5-0103), and two compounds categorized as ‘miscellaneous’ as they did not have the same function as other epigenetic modifiers in the library. Out of the two compounds categorized as miscellaneous, compound Cas # 160003-66-7 (iniparib), has an unknown function and compound Cas # 1207113-88-9 (CCG-100602) is an inhibitor of Rho pathway-mediated signaling [40]. The positive control, ruxolitinib, also reduced intracellular IL-6 by greater than fourfold as previously shown [33]. As HDACi accounted for six out of nine ‘hits’, HDACi were selected for further validation *ex vivo* on cells from aviremic individuals living with HIV. We examined SBHA and panobinostat, two of the top HDACi to reduce intracellular IL-6, for their ability to reduce secretion of a broad panel of pro-inflammatory cytokines *ex vivo* (**Figure 2**).

**Figure 1.**
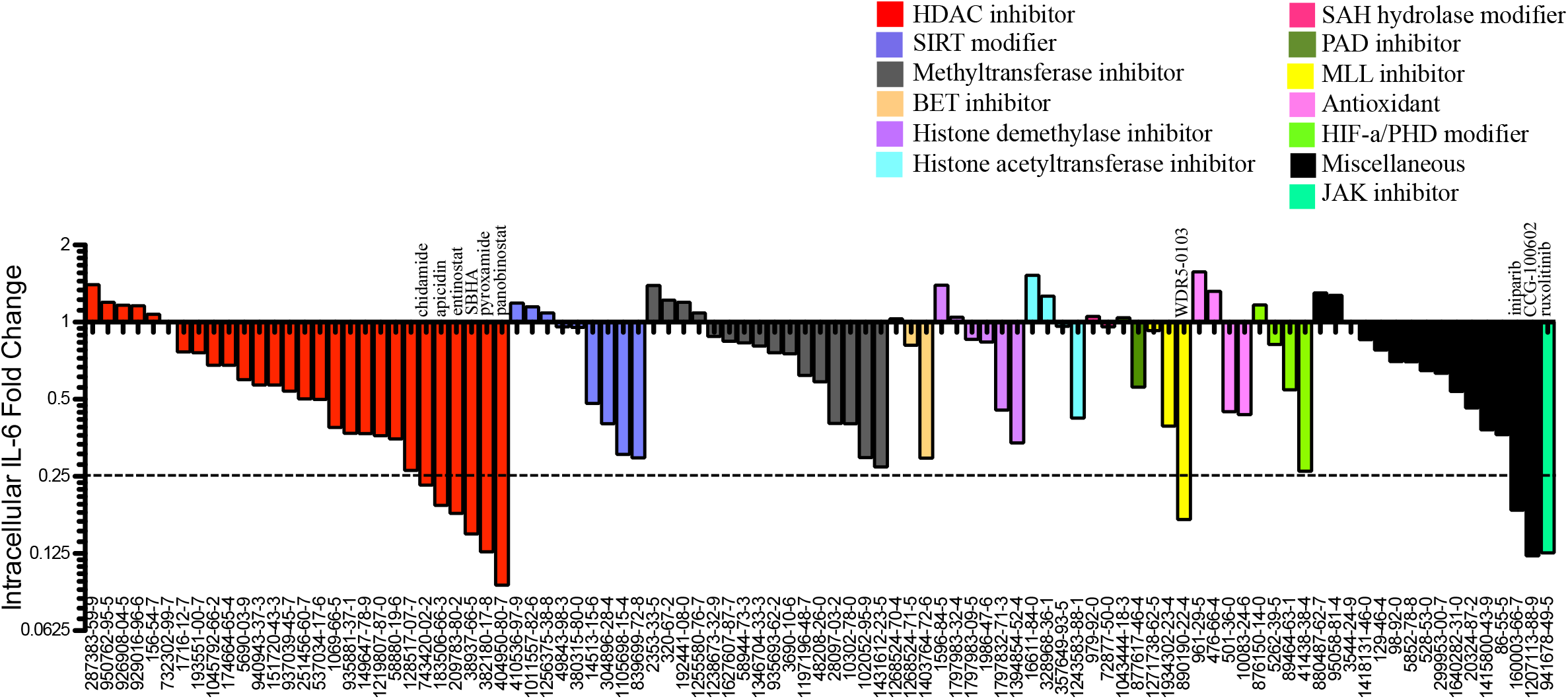
Epigenetic modifiers screening identifying compounds that dampen cytokine production induced by Ingenol-3,20-dibenzoate. Screening of 96 epigenetic modifiers for compounds that reduced intracellular IL-6 induced by IDB in healthy donor PBMCs (n=1) revealed nine epigenetic modifier ‘hits’ that reduce intracellular IL-6 by fourfold or greater (at or below dotted line) when compared to IDB treatment alone. Ruxolitinib was included as a positive control for reduction of intracellular IL-6 when combined with Ingenol-3,20-dibenzoate. Compounds are listed by CAS number and color coded according to categorized function. All nine ‘hits’ were selected for further testing.

**Figure 2A).**
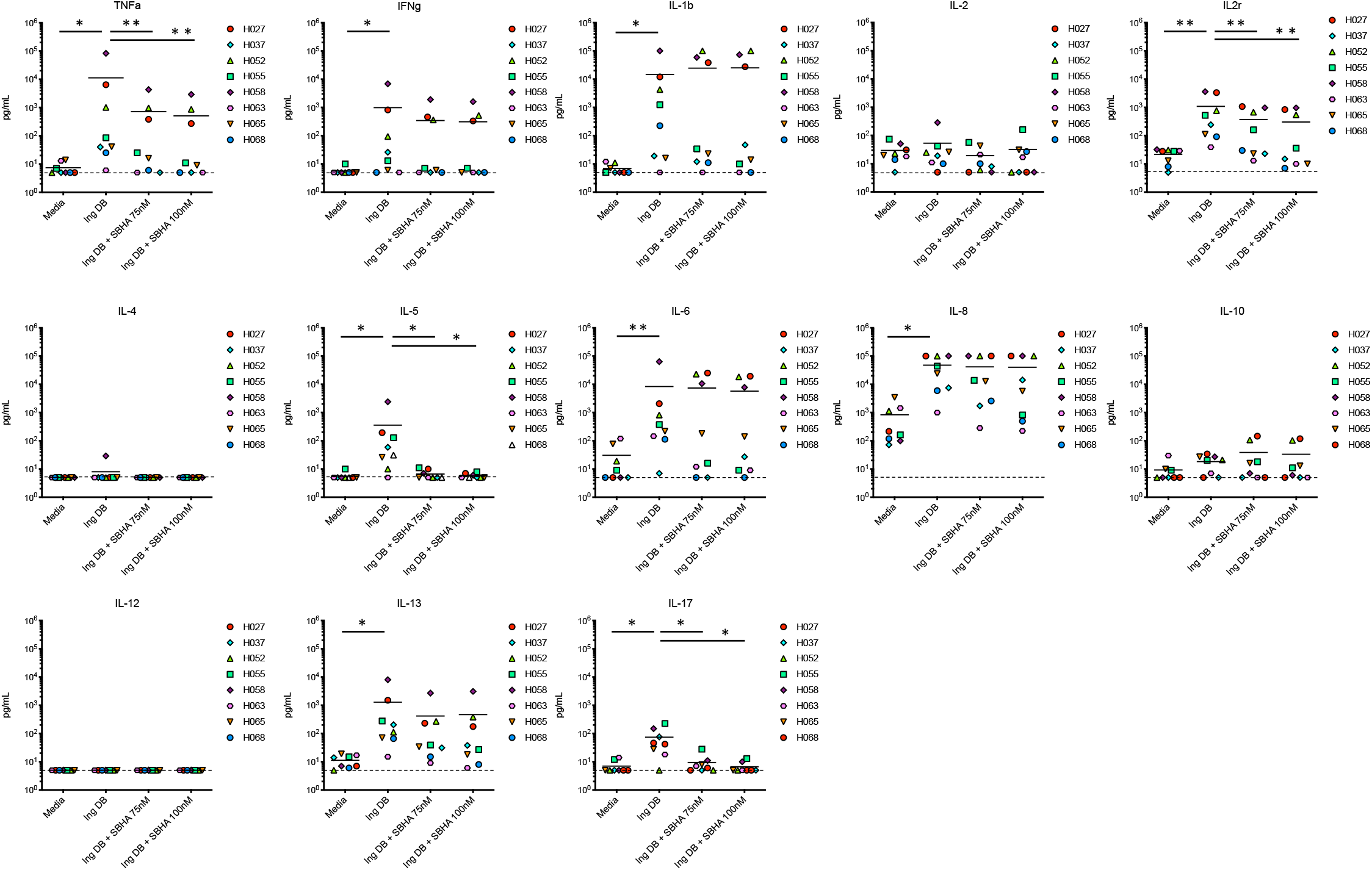
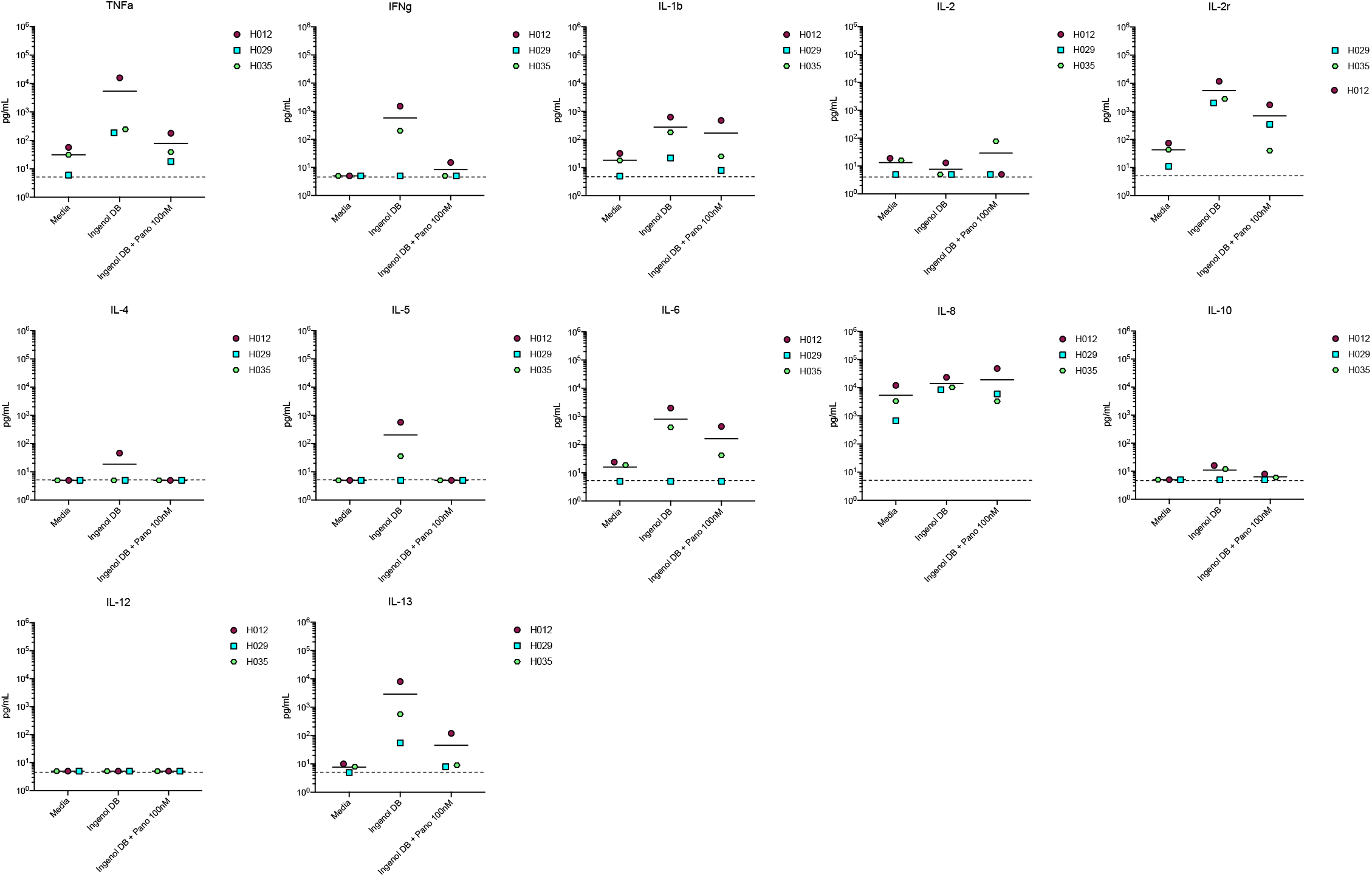
SBHA reduces pro-inflammatory cytokine secretion induced by IDB ex vivo. Solid lines represent the mean change in pro-inflammatory cytokine concentrations in the supernatant of PBMCs isolated from HIV-1 positive aviremic individuals (n = 8) and treated with SBHA (75nM or 100nM) and IDB (100nM) for 72 hours compared to IDB alone. The dotted line represents the baseline for the assay. *P value <0.05; **P value <0.01. 2**B)** Panobinostat has no significant effect on IDB induced pro-inflammatory cytokine production ex vivo. PBMCs from aviremic individual was treated with Panobinostat (100nM) and IDB (100nM). The mean change in pro-inflammatory cytokines is indicated by a solid line and dotted lines indicate the baseline for the assay.

Treatment of aviremic patient PBMCs with IDB significantly increased TNF-α (p = 0.0156), IFN-γ (p = 0.0312), IL-1β (p =0.0156), IL-5 (p = 0.0156), IL-8 (p = 0.0156), IL-13 (p = 0.0156), and IL-17 (p = 0.0156) (**Figure 2A**). The addition of 75nM or 100nM SBHA to 100nM IDB was found to significantly reduce secreted levels of TNF-α (p = 0.0078), IL-5 (p = 0.0156), IL-2r (p = 0.0078), and IL-17 (p = 0.0156) (n = 8) (**Figure 2A**) ex vivo. For IL-13 statistical significance was not achieved (p = 0.0547) upon treatment of cells with IDB and 75nM SBHA. However, the addition of SBHA to IDB stimulation did not diminish the induction of IFN-γ and IL-8, which are significantly induced by IDB treatment. The effect of SBHA on IDB-induced IL-6 and IL-1β levels appeared inconsistent between donors *ex vivo*, with donors clustered into two distinct populations. Three donors (H027, H052, and H058) appeared to cluster above the mean IL-6 level when donor PBMCs were treated with IDB and SBHA at 75nM or 100nM. In the remaining 5 donors (H037, H055, H063, H065 and H068), IL-6 appeared to decrease to near the limit of detection, though this decrease was not statistically significant for the overall population (n = 8) when treated with either 75nM (p =0.0625) or 100nM (p = 0.1250) SBHA. Secreted IL-2, IL-4, IL-10, and IL-12 levels remaining unchanged upon treatment with IDB and the combination therapy. The addition of 100nM panobinostat to 100nM IDB did not significantly reduce secreted levels of any cytokines measured (**Figure 2B**; TNF-α (p = 0.2500), IFN-γ (p = 0.5000), IL-1β (p = 0.2500), IL-2 (p > 0.9999), IL-2r (p = 0.2500), IL-4 (p > 0.9999), IL-5 (p = 0.5000), IL-6 (0.5000), IL-8 (p > 0.9999), IL-10 (0.5000), IL-12 (no difference), IL-13 (p = 0.2500)) (Figure 3B). Secreted IL-17 was not measured.

**Figure 3).**
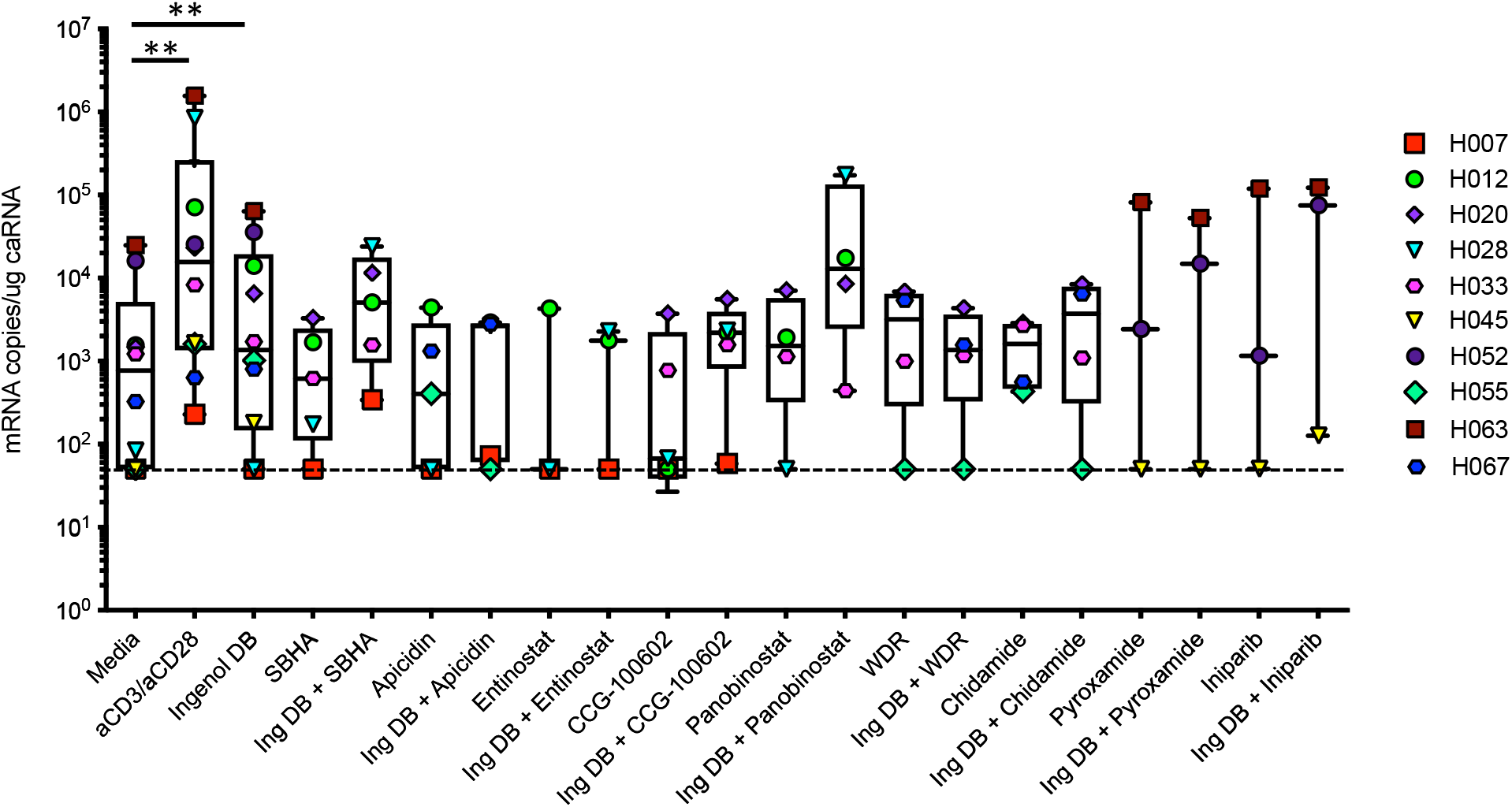
HIV-1 latency reversal induced by epigenetic modifiers alone or in combination with Ingenol-3,20-dibenzoate *ex vivo*. Treatment of rCD4 cells isolated from HIV-1 positive aviremic individuals with Ingenol-3,20-dibenzoate significantly increased viral latency reactivation compared to media alone. The positive control αCD3/αCD28 resulted in viral reactivation in all donors. All nine epigenetic modifiers alone or in combination with Ingenol-3,20-dibenzoate did not increase reactivation significantly compared to Ingenol-3,20-dibenzoate treatment alone. *P value <0.05; ** P value <0.01. The dotted line indicates the limit of detection of the assay (50 copies/ug cell associated RNA (caRNA)).

We then examined the ability of SBHA and panobinostat to reactivate latent HIV-1 when used in combination with IDB *ex vivo* in aviremic patient CD4^+^ T cells. While treatment with IDB or αCD3/αCD28 significantly induced reactivation of the latent reservoir (mean = 12,406 mRNA copies/mL, p value = 0.0078 and mean = 254,182 mRNA copies/mL, p value = 0.0020 respectively, n=10) when compared with media alone, treatment with IDB in combination with SBHA (p = 0.0625) or panobinostat (p = 0.2500) did not significantly reactivate latent HIV-1 (**Figure 3**). Upon testing the remaining epigenetic modifier ‘hits’ identified in our *in vitro* cytokine screen, none of the remaining epigenetic modifiers significantly reactivated latent HIV in patient samples when combined with IDB (apicidin (p = 0.1250), entinostat (p = 0.5000), CCG-100602 (p = 0.0625), WDR5-0103 (p = 0.5000), chidamide (p = 0.5000), pyroxamide (p = >0.9999), iniparib (p = 0.2500)) compared to media alone. Additionally, all nine epigenetic modifiers when used individually failed to reactivate latent HIV-1 to significant levels (**Figure 3**).

## 4. Discussion

In order to eliminate the latent HIV-1 reservoir, LRA therapies must be evaluated based on both their potency and deleterious side effects. PKC agonists are among the most potent LRAs that have emerged across multiple models of HIV-1 latency [29, 30]. However, PKC agonists are also known to induce T cell activation [32] and secretion of pro-inflammatory cytokines [33, 41]. Previous reports have demonstrated that LRAs can be combined with a second pharmacological agent to suppress deleterious pro-inflammatory cytokine secretion induced upon treatment with PKC agonists [33] and TCR stimulation [34]. While HDAC inhibitors such as suberanilohydroxamic acid (SAHA) [35, 37], Trichostatin A [36], and Nicotinamide [36] have been shown to reduce pro-inflammatory cytokine secretion in other disease contexts [35, 36], we here, for the first time, have examined the ability of HDACi and other epigenetic modifiers to do so in the context of HIV-1.

We found that SBHA, an HDAC1-3 inhibitor, significantly reduced Ingenol-3,20-dibenzoate (IDB) induced secreted TNF-α, IL2-r, IL-5, and IL-17 in PBMCs from aviremic individuals living with HIV-1 *ex vivo*. While SBHA has been previously described to induce cell cycle arrest [42], apoptosis [42-44], cell differentiation [45], and has been shown to decrease Herpes simplex virus type 1 viral genome expression *in vitro* [46], SBHA has not been described to inhibit the production of pro-inflammatory cytokines. However, HDAC1-3 inhibition is a known mechanism of inhibiting the production of pro-inflammatory cytokines [47-50]. HDAC1- and HDAC3-independent knock-down have been previously shown to reduced LPS-induced IL-6 and TNF-α in murine microglia [47] and IL-1 pro-inflammatory cytokine induction in HEK293IL-1R and mouse embryonic fibroblasts [50]. Additionally, other HDAC1-3 inhibitors such as entinostat have been shown to reduce the production of pro-inflammatory cytokines IL-1β and IL-6 in a rat collagen-induced arthritis model [48] and inflammatory gene mRNA levels for IL-1β, IL-17, and IFN-γ in an experimental autoimmune neuritis rat model [49]. In this study we showed that entinostat reduced intracellular IL-6 production but to a lower degree than SBHA, panobinostat, and pyroxamide and was therefore not studied further. However, it does strengthen the case that HDAC1-3 inhibition is a candidate pathway for inhibiting pro-inflammatory cytokines induced by PKC stimulation during HIV-1 latency reversal, as we demonstrated by combining known HDAC1-3 inhibitor SBHA with IDB to reduce the production of PKC-induced TNF-α, IL-17, IL-2r, and IL-5 *ex vivo*. While SBHA reduced PKC-induced pro-inflammatory cytokines, panobinostat, a pan-HDAC inhibitor, did not significantly reduce pro-inflammatory cytokines *ex vivo*. Panobinostat has been previously shown to significantly reduce IL-6 in a Hodgkin’s Lymphoma phase II clinical trial [51] and IFN-γ secretion in a lymphocyte-target cell killing assay [52]. Other pan-HDACi such as TSA have been reported to reduce IL-6 [36] and TNF-α [36, 53] secretion in LPS-stimulated macrophages, although conflicting studies have shown that TSA has no effect on other pro-inflammatory genes such as IL-1β and IL-10 [53]. It is unclear why panobinostat did not reduce PKC-induced pro-inflammatory cytokine production *ex vivo*.

In addition to reducing PKC-induced pro-inflammatory cytokines, we aimed to identify an epigenetic modifier which could be used in combination with IDB without reducing latency reversal. HDACi such as SAHA [24, 54, 55] and panobinostat [56] have been previously shown to reactivate latent HIV-1in clinical trials. However, *in vitro* primary cell models of HIV-1 latency and *ex vivo* aviremic patient cell studies have produced conflicting results as to the ability of panobinostat and SAHA to reactivate latent HIV-1 [28, 57-59]{Spina, 2013, 24385908}. In this study, panobinostat did not significant reactivate latent HIV-1 in our *ex vivo* model, as has been seen previously [57]. SAHA was not evaluated as a LRA in this study as it did not significantly reduce PKC-induced intracellular IL-6 in our *in vitro* screen.

## 5. Conclusions

The combination therapy of SBHA and IDB significantly reduced PKC-induced pro-inflammatory cytokines TNF-α, IL-17, IL-2r, and IL-5 suggesting this combination therapy may be an effective strategy to inhibit pro-inflammatory cytokine induction in the context of HIV-1. However, reduction of pro-inflammatory cytokines by SBHA prevented significant latency reversal by IDB suggesting that combining SBHA with IDB would not be an efficient strategy to reactivate latent HIV-1.

## Acknowledgements

The authors express their sincere gratitude to the study participants for their continued participation in this work and ongoing translational research. This work was supported in part by the National Institute of Allergy and Infectious Diseases (NIAID) grants AI122377-01 and AI143567-02 (VP), NIAID Ruth L. Kirschstein National Research Service Award T32AI055434-11 (ETL), NIAID grant R21AI124823-02 (LRB), and the Doris Duke Charitable Foundation Clinical Scientist Development Award CSDA201612 (AMS). The research reported in this publication was supported by the National Center for Advancing Translational Sciences of the National Institutes of Health under Award Number UL1TR002538. The content is solely the responsibility of the authors and does not necessarily represent the official views of the National Institutes of Health.

## Ethics approval

Heathy donors and HIV-1 positive aviremic individuals on ART were recruited for phlebotomy according to Institutional Review Board (IRB) protocols 67637 (approved May 31, 2017) and 58246 (approved Jan 4, 2017) at the University of Utah.

